# The effect of weak clonal interference on average fitness trajectories in the presence of macroscopic epistasis

**DOI:** 10.1101/2021.08.12.456174

**Authors:** Yipei Guo, Ariel Amir

## Abstract

Adaptation dynamics on fitness landscapes is often studied theoretically in the strong-selection, weak-mutation (SSWM) regime. However, in a large population, multiple beneficial mutants can emerge before any of them fixes in the population. Competition between mutants is known as clonal interference, and how it affects the form of long-term fitness trajectories in the presence of epistasis is an open question. Here, by considering how changes in fixation probabilities arising from weak clonal interference affect the dynamics of adaptation on fitness-parameterized landscapes, we find that the change in the form of fitness trajectory arises only through changes in the supply of beneficial mutations (or equivalently, the beneficial mutation rate). Furthermore, a depletion of beneficial mutations as a population climbs up the fitness landscape can speed up the functional form of the fitness trajectory, while an enhancement of the beneficial mutation rate does the opposite of slowing down the form of the dynamics. Our findings suggest that by carrying out evolution experiments in both regimes (with and without clonal interference), one could potentially distinguish the different sources of macroscopic epistasis (fitness effect of mutations vs. change in fraction of beneficial mutations).

## INTRODUCTION

For unicellular organisms, as individuals in an asexual population gain mutations and increase in fitness, how their average fitness increases over evolutionary timescales (which we refer to as the average fitness trajectory *F* (*t*)) depends on the distribution *ρ*(*s*) of fitness effects *s* of potential new mutations (i.e. mutations that can potentially occur). This distribution may change as a population evolves, an effect known as macroscopic epistasis (Fig. 1a) [1]. This could come about, for example, when the fitness effects of mutations depend on the state of other genes or the presence of other mutations [2–4]. The effect of such changes in *ρ*(*s*) on fitness trajectories has been studied in the context of fitness-parameterized landscapes, where *ρ*(*s|x*) is assumed to depend only on the fitness *x* of the cell [1, 5].

**FIG. 1.**
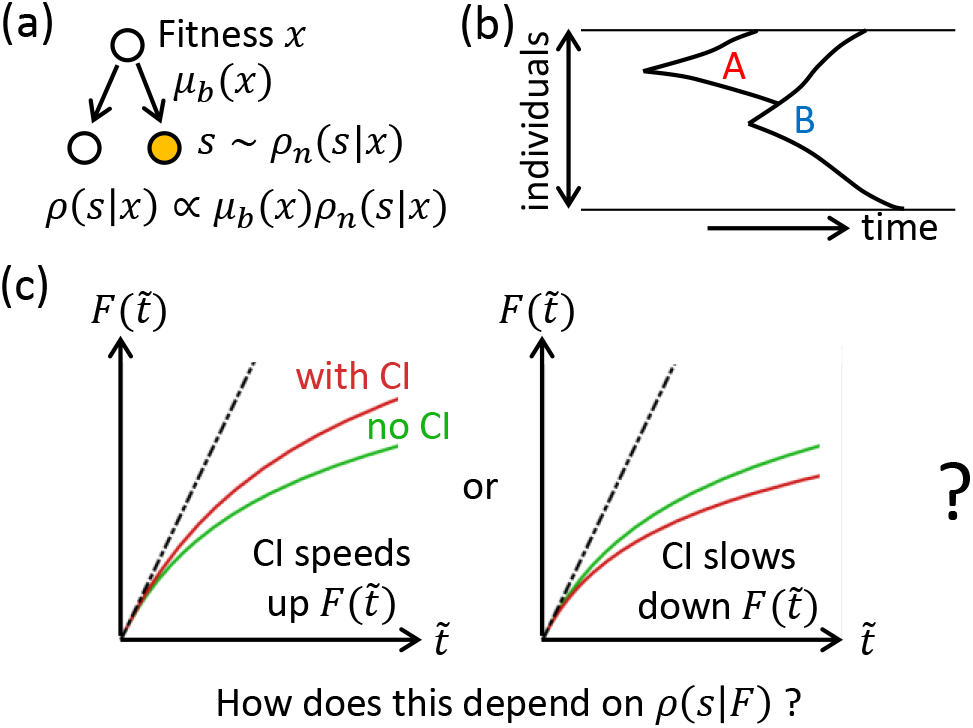
Schematic of the model and the question-of-interest. (a) Whenever a cell divides, there is a probability *μ_b_*(*x*) of it gaining a beneficial mutation which can depend on the fitness *x* of the cell. When a beneficial mutation occurs, we assume that the selection coefficient *s* of the mutation is drawn from some normalized distribution *ρn*(*s|x*) which can also depend on *x*. The overall distribution *ρ*(*s|x*) of beneficial selection coefficients when a mutation occurs specifies the fitness-parameterized landscape, or equivalently, the form of macroscopic epistasis. (b) Clonal interference refers to the phenomenon where a mutant (‘A’, already growing exponentially in size) is out-competed by a new mutant (‘B’) with a higher selective advantage. (c) The functional forms of the average fitness trajectories *F* (*t*) are compared after rescaling time such that the initial adaptation speeds are identical. We ask whether clonal interference speeds up (left) or slows down (right) the functional form of the average fitness trajectory, and how this depends on *ρ*(*s|F*).

Within this framework, adaptation dynamics observed experimentally can be used to infer features of the underlying fitness landscape or equivalently, the type of epistasis present in the system [1, 6, 7]. The shape of fitness trajectories (which we characterize using its functional form) is typically studied theoretically in the strong-selection, weak-mutation (SSWM) regime, in which the time for beneficial mutations to fix is much shorter than the time for successful beneficial mutations to emerge [8]. However, in large populations, multiple beneficial mutants can emerge before any of them fixes. Competition between beneficial mutants in different lineages is known as clonal interference (Fig. 1b) and has been shown to reduce fixation probabilities [9, 10]. There have also been suggestions that clonal interference slows down the form (i.e. shape) of fitness trajectories [6, 11] (here, we are interested in the functional forms of the fitness trajectories, and therefore the trajectories are compared after scaling time such that they have identical initial slopes —we consider a trajectory to be slower than another if it has a slower fitness increase after this re-scaling, Fig. 1c). In particular, the slow fitness trajectory observed in Lenski’s long-term evolution experiment (where the fitness appears to continue increasing over a long period of time without reaching any plateau) was previously attributed to both diminishing returns epistasis and clonal interference [6]. However, the relative contributions from these two factors have not been explored, and it is not clear if it is generally the case that clonal interference slows down the form of fitness trajectories in the presence of epistasis.

Here, by considering how changes in fixation probabilities arising from weak clonal interference affect the dynamics of adaptation on fitness-parameterized landscapes, we find that the change in the form of fitness trajectory due to clonal interference arises only through changes in the supply of beneficial mutations (or equivalently, the beneficial mutation rate), independent of any changes in the average fitness effect of beneficial mutations. This implies that, contrary to what has been argued in the literature, as long as the fraction of beneficial mutations stays the same, the functional form of the fitness trajectory will be the same with or without clonal interference, even if the mean fitness effect of beneficial mutations decreases over time (i.e. diminishing returns). Furthermore, a depletion of beneficial mutations as a population climbs up the fitness landscape can speed up the functional form of the fitness trajectory (while an enhancement of the beneficial mutation rate slows it down). These findings suggest that by carrying out evolution experiments in both regimes (with and without clonal interference), one could potentially distinguish the different sources of macroscopic epistasis (change in fitness effects of mutations vs. change in the fraction of beneficial mutations).

## RESULTS

We consider the Moran process, where a population has a constant size *N*, and each cell divides at some rate which we call the fitness of the cell. Whenever a cell divides, a random cell is simultaneously removed from the population. Experimentally, this corresponds, approximately, to growing cells in a continuous culture (e.g. in a turbidostat) [12, 13]. During each division event, there is some probability that the daughter cell gains a new mutation (Fig. 1a). We assume that the fitness effects of deleterious mutations are typically much larger than those of beneficial ones, such that beneficial mutations rarely compensate deleterious mutations, and hence deleterious mutations cannot fix. The adaptation of the population is therefore driven by the probability *μ_b_* of gaining a beneficial mutation during division, which is a dimensionless quantity that we will refer to as the beneficial mutation rate.

Inspired by previous theoretical studies [14, 15], we assume an exponential distribution for the fitness effects of beneficial mutations. Such a distribution also seems to have some experimental support [16–18]. Nevertheless, there are other data that suggest otherwise [19, 20], and we show in SI sections III, IV that our main conclusions hold for a more general class of distributions. Generalizing the approach of Ref. [5], given the current fitness of a cell *x*, the distribution of its beneficial mutant fitness values *y* > *x* is assumed to be given by:

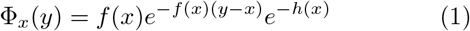

where 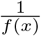 represents how the mean fitness effects (*y* − *x*) of mutations vary with current fitness, and *h*(*x*) governs how the fraction of beneficial mutations changes with the current fitness of the cell. The beneficial mutation rate is then given by *μ_b_*(*x*) = *μ*_b0_*e*^−*h*(*x*)^, with *μ*_*b*0_ being the initial beneficial mutation rate.

For any Φ_*x*_(*y*), the corresponding distribution of fitness selection coefficients 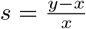 is then given by

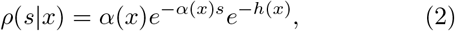

where *α*(*x*) = *xf* (*x*) = ⟨*s*⟩ ^−1^ is the inverse of the mean selection coefficient of beneficial mutations. Within this model, macroscopic epistasis can act through changing the mean of beneficial mutation effects as the population increases in fitness, through changing the availability of new beneficial mutations, or a combination of both.

This framework reduces back to other previously known models for specific forms of *f* (*x*) (or equivalently, *α*(*x*)) and *h*(*x*). For example, the House of cards (HoC)/Uncorrelated fitness landscape is the case where Φ_*x*_(*y*) is independent of *x* and corresponds to *f* (*x*) ~ 1 and *h*(*x*) ~ *x* [5]. This implies that while the absolute mean fitness increase conferred by mutations stay the same, the availability of beneficial mutations decreases exponentially as fitness increases. Similarly, the nonepistatic (NEPI) fitness landscape is the case where the distribution of fitness effects of mutations is independent of genotype, i.e. Φ*_x_*(*y*) is only a function of *y* − *x* [5]), which corresponds to having *f*(*x*) ~ 1 and *h*(*x*) = 0. The Stairway to heaven (STH) fitness landscape [5] is the case where *ρ*(*s*) is independent of fitness, and corresponds to having *α*(*x*) ~ 1 while *h*(*x*) = 0. The diminishing returns epistasis model adopted by Wiser et al. in Ref. [6] can also be mapped to this fitness-parameterized framework with *α*(*x*) ~ *x*^*g*^ for *g* ≫ 1 and *h*(*x*) = 0.

We assume that the beneficial mutation rate is not too large such that the population is typically in a monomorphic state. When a new beneficial mutation with fitness effect *s* emerges, it fixes with some probability *P_f_* (*s*). The average fitness trajectory *F* (*t*) on such fitness-parametrized landscapes is then approximately given by [5]:

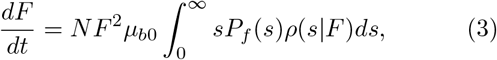

where time *t* here refers to real time (i.e., the actual amount of time that has passed in a continuous culture), in contrast to being the number of generations often considered in other studies [5, 6].

In the SSWM regime (*Nμ_b_*log(*Ns*) ≪ 1 [8]), *P_f_* (*s*) = *π*(*s*) ≈ *s*. Substituting *ρ*(*s|F*) from Eqn. 2 and defining 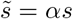 (the selection coefficient of a mutation relative to the average selection coefficient), the dynamics of fitness in this regime are given by:

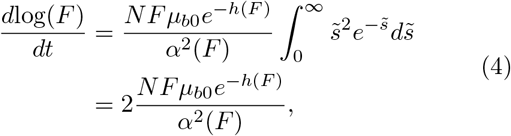

where the functional form of the fitness trajectory depends on the type of epistasis specified through *α*(*F*) and *h*(*F*). For example, the HOC landscape (*h*(*F*) ~ *F*, *α*(*F*) ~ *F*) gives *F* (*t*) ~ log(*t*), the NEPI landscape (*h*(*F*) = 0, *α*(*F*) ~ *F*) gives *F* ~ (*t*) *t*, and the STH landscape (*α*(*F*) ~ 1, *h*(*F*) = 0) gives a function *F* (*t*) that increases faster than linearly. The diminishing re-turns model by Wiser et al. (*α*(*F*) ~ *F^g^*, *h*(*F*) = 0) gives a power law trajectory with exponent 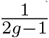.

When a population is sufficiently large, after a mutation has escaped loss via genetic drift and the number of mutants starts to increase deterministically, there is some probability that another new successful mutation emerges in the resident population while the first mutant is still in the process of taking over the population. Following the approach in Ref. [9], for a mutant that emerges in a clonal population containing individuals with fitness *F*, the probability of it fixing *P_f_* (*s*) will then be the probability that it escapes loss via genetic drift (and would go on to fix in the absence of any interference) *π*(*s*), and not outcompeted by any other mutation that could potentially emerge in the background population while the first mutant population is growing in size:

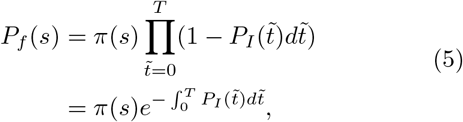

where 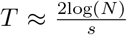 is the average number of generations it takes for a mutant to fix without any interference (SI section I). 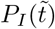 is the probability that a successful interfering mutation emerges at time 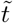 after the first mutant sub-population starts growing in size, and is given by the probability that the resident population gains another more beneficial mutation which subsequently goes on to fix:

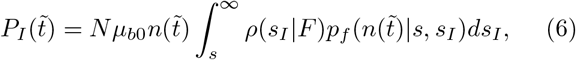

where 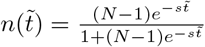 is the fraction of resident cells (SI section I), *ρ*(*s_I_*|*F*) is the distribution of fitness effects of mutations that can arise in a cell with fitness *F* (Eqn. 2), and 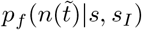 is the fixation probability of the interfering mutant with fitness effect *s_I_*. Intuitively, we expect *p_f_*(*n|s*, *s_I_*) to increase with *n* since the closer the first mutant is to fixing (i.e. lower *n*), the harder it is for the new mutant to outcompete the original mutant. In particular, one might also expect the probability of the interfering mutant fixing in this mixed population to be the same as that in a clonal population with the same average population growth rate (i.e. all cells grow at this rate). We will show that this is indeed true for all *n* when the growth rate of the mutant is much larger than that of the population-averaged growth rate (but not in general).

Assuming that clonal interference is weak such that at most one successful competing mutation occurs during the time a mutant is trying to fix (i.e. no further mutations occur after the emergence of the second mutant), the extinction probability *p*_*ext*_(*n*) of the new (interfering) mutant (in the large *N* limit) can be found by considering the different events that can happen (different cell types dividing and leaving the population) in the next time step and requiring all remaining mutants to go extinct (SI section II). The fixation probability *p*_*f*_(*n*) = 1 − *p*_*ext*_(*n*) is then found to be given by the solution to the following equation (SI section II):

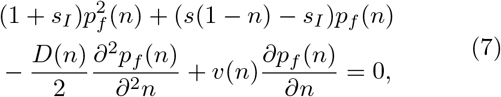

where 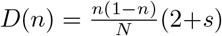 and *v*(*n*) = *n*(1 − *n*)*s*, and the boundary conditions are given by *p*_*f*_(0) = (*s*_*I*_ − *s*)/(1 + *s*_*I*_) and *p*_*f*_ (1) = *s*_*I*_/(1 + *s*_*I*_), which are also the fixation probabilities of a mutant in a clonal population within the Moran process [13, 21].

When the first two terms in Eqn. 7 dominate (either when *n* is close to 0 or 1, or when *s*_*I*_ ≫ *s*), we recover the intuitive result that the fixation probability is the selective advantage of the interfering mutant over that of the population average:

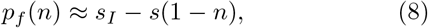

where we have also taken the limit where fitness effects are generally small *s*, *s*_*I*_ ≪ 1. For simplicity and concreteness, we will assume this specific form for *p_f_* (*n*) in the rest of the paper, but we argue in SI section III that adopting the more general solution of Eqn. 7 does not affect our main conclusions.

Within this approach, we have considered the effect of potential interfering mutants on the fixation probability of a mutant that initially arose in a clonal population. This implicitly assumes that clonal interference is sufficiently weak that most of the time a mutation emerges when the population is dominated by a single subgroup. We elaborate on this assumption more extensively in the Discussion section.

The dynamics of fitness in the presence of weak clonal interference is therefore given by (using Eqns. 3, 5, 6, 8, see SI section III):

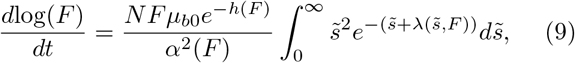

where, in the limit log(*N*) ≫ 1,

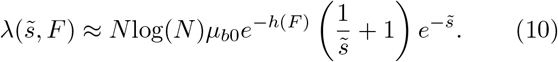

If the beneficial mutation rate does not depend on fitness (*h*(*F*) = 0), we recover the same expression for 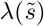 derived by Gerrish and Lenski in Ref. [9].

In the following section, we will explore the effect of clonal interference on the functional form (i.e. shape) of the average fitness trajectory by comparing the dynamics with (Eqn. 9) and without (Eqn. 4) clonal interference. When comparing the numerical solutions of Eqns. 4 and 9, we scale time in the case with clonal interference by a factor such that 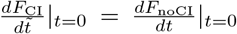 (since we are interested in comparing functional forms rather than the absolute time dynamics).

To verify our findings, we also carried out full Gillespie simulations of the Moran process, where we keep track of the fitness values of all cells, with the probability of a cell dividing proportional to its fitness (see SI section V for simulation details). When a mutation event occurs during division, the fitness effect of the mutation is drawn from a distribution based on the fitness of the dividing cell. Within each simulation, the mean population fitness at any time is obtained from an average over all cells in the population.

### Change in functional form of fitness trajectory only depends on how beneficial mutation rate changes

Since any additional dependence on *F* due to the clonal interference term comes only from *h*(*F*) (Eqns. 9, 10), we expect weak clonal interference to only affect the functional form of the fitness trajectory if the beneficial mutation rate *μ_b_* = *μ*_*b*0_*e*^−*h*(*F*)^ changes with fitness (Fig. 2). In other words, if *h*(*F*) ~ *O*(1) (such as in the case of the diminishing returns model, the NEPI, and the STH landscapes), even if there is epistasis that acts through *α*(*F*), clonal interference does not change the functional form of the trajectory (Fig. 2a). This is the case because as long as the availability of beneficial mutations does not change, the presence of clonal interference reduces the rate of fitness increase by the same amount at all times.

**FIG. 2.**
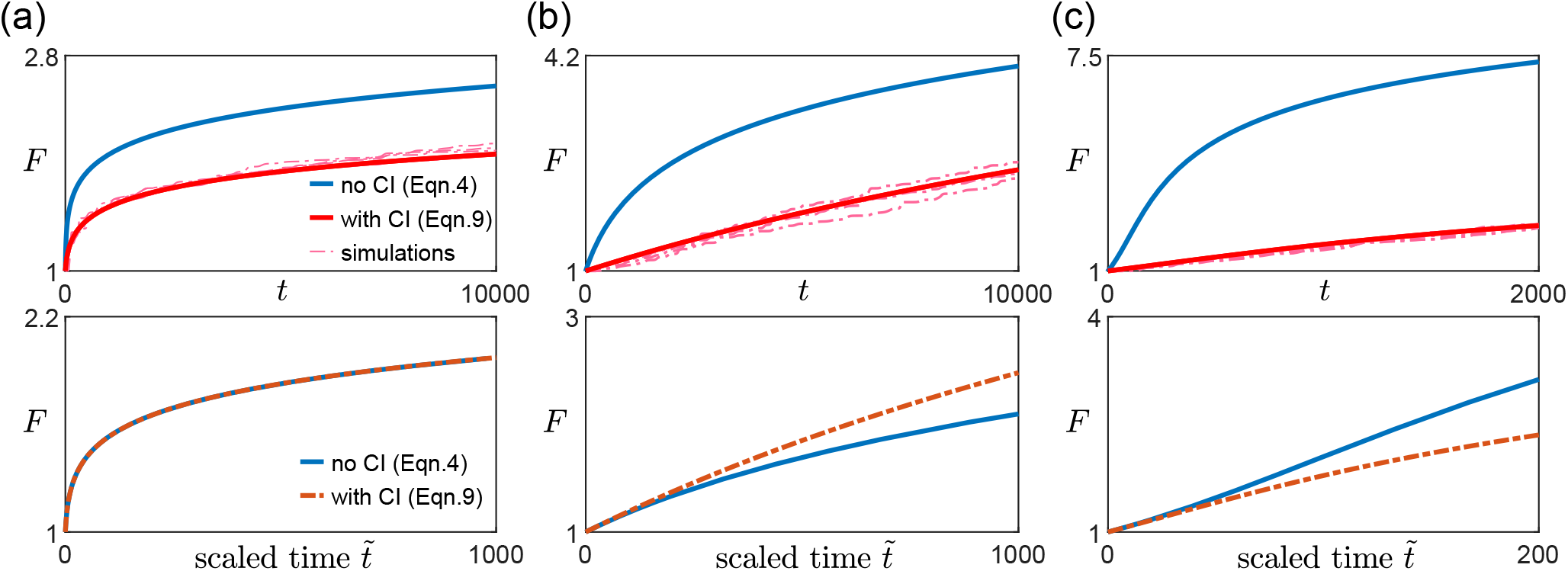
Effect of clonal interference on fitness trajectory for different types of macroscopic epistasis. (a) Within the diminishing returns model where the mean fitness effect of beneficial mutations decreases with fitness but beneficial mutation rate stays the same, the functional form of the fitness trajectory stays the same. The upper panels show the fitness trajectories as a function of actual time *t* with (red) and without (blue) clonal interference. The solid lines are are obtained from solving Eqns. 4 and 9, while the dashed lines (pink) are individual replicates from simulations. The lower panels compare trajectories (blue and red solid lines in the upper panel) after rescaling time 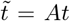 for the case with clonal interference, with *A* being a constant chosen such that both trajectories have the same derivative at 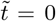 [Landscape parameters: *α*(*F*) = 20*F* ^5^, *μ*_*b*_ = *μ*_*b*0_] (b) With the House-of-Cards/uncorrelated landscape where the mean fitness effects of beneficial mutations stays the same but the beneficial mutation rate decreases with fitness, the presence of clonal interference speeds up the functional form of the fitness trajectory. [Landscape parameters: *α*(*F*) = 100*F*, *μ*_*b*_ = *μ*_*b*0_*e*^−(*F* − 1)^] (c) With a mutation-releasing landscape where beneficial mutation rate increases with fitness, the presence of clonal interference slows down the functional form of the fitness trajectory.[Landscape parameters: *α*(*F*) = 50*e*^*F* − 1^, *μ*_*b*_ = *μ*_*b*0_*e*^*F*−1^.] [Other parameters in (a)-(c): *N* = 10^6^, *μ*_*b*0_ = 1 *·* 10^−5^.]

If *h*(*F*) depends on *F*, whether the form of the trajectory is sped up or slowed down by clonal interference depends on whether *h*(*F*) increases or decreases with *F*. In many commonly used and widely-studied models of fitness landscapes, *h*(*F*) increases with *F*, i.e., the beneficial mutation rate decreases with fitness. For example, in the HoC landscape, a new fitness value is drawn from the same distribution whenever a mutation occurs [5, 22], implying that as fitness increases, the probability of drawing a higher value of fitness (for the mutant) decreases. In Fisher’s geometric model [23], the state of the cell lies in a multi-dimensional phenotypic space, with a certain point being the optimal phenotype, such that the fitness is specified by how close the state of the cell is to the optimal state. Mutations then correspond to drawing random vectors in the space. Within this model, the probability of a mutation providing an improvement (i.e. bringing the population closer to the optimal state) also decreases with fitness [23–25]. In these cases where *h*(*F*) increases with *F*, clonal interference speeds up the functional form of the trajectory (Fig. 2b). Intuitively, this occurs because as fitness increases, the effect of clonal interference is reduced since there are fewer potential beneficial interfering mutations.

Similarly, in the opposite case where *h*(*F*) decreases with *F*, there are more potential beneficial mutations as the population climbs up the fitness landscape (one could imagine this happening if gaining certain mutations opens up more beneficial mutational paths). In this case, interfering mutations become more common as the population evolves and the form of the trajectory is slowed down by such an effect (Fig. 2c).

Besides the average fitness trajectory, we find that these general arguments and results also hold for the average substitution trajectory, i.e., how the average number of fixed mutations increases over time (SI section VI).

### Experimental protocol for determining how beneficial mutation rate changes

To determine whether and how the availability of beneficial mutations changes over time (with fitness), one can carry out two or more sets of long-term evolution experiments at different population sizes or mutation rates [26] spanning both the SSWM regime and the regime with weak clonal interference (Fig. 3a). *N* can be varied, for example, by carrying out experiments in a turbidostat at different optical densities, while potential ways of varying *μ* include changing the expression levels of DNA repair enzymes, such as mutH [27] and Ada [28] in *E. coli*, and inducing mutagenesis using UV radiation [29]. The average fitness trajectory for a given *N* and *μ* can then be obtained by averaging over the replicates in the corresponding set of experiments (Fig. 3b). In general, the initial rate of fitness increase will differ between sets of experiments (Fig. 3b). Calculating this initial rate and scaling time for each of these sets such that all scaled fitness trajectories have the same initial derivative (Fig. 3c-e), one can then infer how *μ_b_* changes. In particular, if the different trajectories overlap (or are statistically indistinguishable), it would suggest that *μ_b_* remains approximately constant (Fig. 3c). If instead the scaled trajectories speed up (slow down) as *N* or the overall mutation rate is increased, it would suggest that *μ_b_* decreases (increases) as fitness increases over time (Fig. 3d, e).

**FIG. 3.**
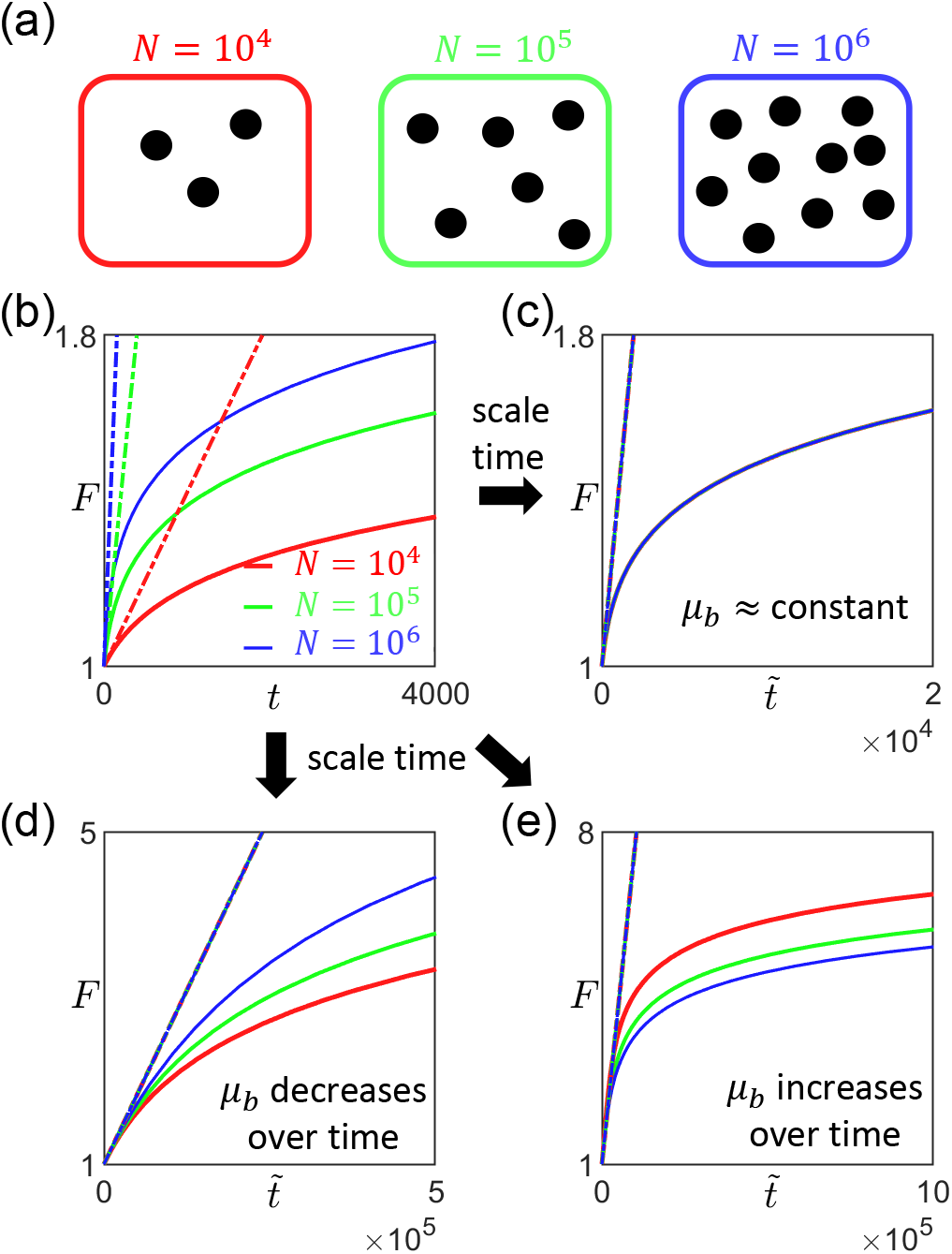
Proposal for experimental protocol to determine how the availability of beneficial mutation changes as fitness increases. (a) The first step is to carry out multiple sets of long-term evolution experiments at different population sizes or mutation rates that span both the SSWM regime and the regime with weak clonal interference. For example, if the initial strain has estimated beneficial mutation rate *μ*_*b*0_ ≈ 10^−5^, one could carry out experiments with *N* = 10^4^, *N* = 10^5^, and *N* = 10^6^. (b) For each set of experiment (with a given *N*), one can then then obtain the average fitness trajectory by averaging over multiple replicates in each set (solid lines), and calculate the corresponding initial rate of fitness increase (dashed lines). (c-e) By scaling time 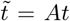 such that the average trajectories have the same initial derivative, one can then infer how the beneficial mutation rate changes over time. If the scaled trajectories approximately overlap for all values of *N*, it would suggest that *μ_b_* remains approximately constant. (d) If the scaled trajectories speed up as *N* is increased, it would suggest that *μ_b_* decreases as fitness increases over time. (e) If the scaled trajectories slow down as *N* is increased, it would suggest that *μ_b_* increases as fitness increases over time.

## DISCUSSION

In the SSWM regime, different types of epistasis have been known to give very similar fitness trajectories. For example, both the HoC landscape (where the mean fitness effect stays the same but beneficial mutations deplete exponentially with fitness) and diminishing returns epistasis with constant beneficial mutation rates can give rise to a slow, approximately logarithmic fitness trajectory that does not seem to approach any plateau. Our results suggest that one way of distinguishing whether it is the average fitness effects of potential mutations or the average fraction of available mutations (or both) that is changing over time is to carry out the same set of evolution experiments with two or more different population sizes or mutation rates [26] (in both the SSWM regime and the regime with weak clonal interference), and compare their fitness trajectories.

Although we have only considered adaptation dynamics on fitness-parameterized landscapes, it is possible for such macroscopic epistasis to arise from a microscopic model of fitness landscape that explicitly takes into account interactions between genes (i.e. microscopic epistasis) [11, 30]. Furthermore, in many of these microscopic models, the rate of beneficial mutations decreases as the population climbs up the fitness landscape. In fact, such depletion of beneficial mutations also occurs when one models the genome as a finite sequence of sites, each having its own independent contribution to fitness (i.e. no microscopic epistasis, in which case *F*(*t*) approaches a maximum *F*_*max*_ according to a power law *F*_*max*_ − *F*(*t*) ~ 1/*t*^2^ in the SSWM regime if the independent contributions to fitness follows an exponential distribution) [1]. Our results therefore suggest that clonal interference can speed up the functional form of fitness trajectories even in these microscopic models.

Even though we have neglected the accumulation of deleterious mutations, if the selection coefficients of deleterious mutations are typically much larger than that of beneficial ones, one might consider the effect of deleterious mutations as decreasing the fraction of cells that can gain successful beneficial mutations [23, 31]. When the deleterious mutation rate is much higher than that of beneficial mutations, the system can be thought of as being in a quasi-equilibrium state (mutation-selection balance), with the fraction of cells having the highest fitness (i.e. free of deleterious mutations) being 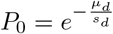 where *μ_d_* is the deleterious mutation rate, and *s_d_* is the harmonic mean of the distribution of fitness effects of deleterious mutations [31]. In the presence of macroscopic epistasis, *μ_d_*, *s_d_*, and hence *P*_0_ can potentially depend on the fitness of the cell. Since successful interfering mutations can only arise in the fraction of resident population that is free of deleterious mutations, there would be an additional factor of *P*_0_(*F*) in the expression for 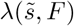 (Eqn. 10), which could further affect the shape of *F* (*t*) (Eqn. 9) in the presence of clonal interference (relative to that in the SSWM regime). Nevertheless, if *s_d_* is independent of *F*, since *μ*_*d*_(*F*) = *μ* − *μ*_*b*_(*F*) (with *μ* being the constant probability of gaining a mutation per division), *P*_0_(*F*) changes (i.e. increases or decreases) with *F* in the same way as *μ*_*b*_(*F*). This implies that the effect of *P*_0_(*F*) in terms of slowing or speeding up the shape of the trajectory is the same as that of *μ_b_*(*F*). In general, *s_d_* could vary with *F* and this could potentially be inferred by comparing fitness trajectories obtained with different *μ* (since this would change the contribution from *P*_0_).

In the regime where the time taken for a mutant to fix is at least comparable to the time for new successful mutations to emerge, besides interference from other beneficial mutations that emerge in the resident population, the original mutant population can also gain additional mutations. The fixation probabilities of these new mutations are enhanced since they are accumulated in cells that already have a fitness advantage in the population. This effect could be important when there are a large number of different subgroups (i.e. genotypes) in the population, such that the average fixation probability of a mutation (of a certain fitness effect) should also be an average over the different background fitness values the mutation can occur in [32]. In general, not accounting for multiple mutations would underestimate the fixation probabilities of small effect mutations since these benefit most from hitch-hiking [33]. However, these small effect mutations also contribute the least amount to fitness, which may explain why taking into account clonal interference alone seems to provide a good agreement with simulations (Fig. 2). Nevertheless, how including the effect of multiple mutations would affect the functional form of long-term fitness trajectories in the presence of epistasis, and how our results extend to the regime of strong clonal interference, are interesting questions that we leave for future work. Another potential direction for future studies is to take into account the interactions between clonal interference and horizontal gene transfer [34] (and how these would affect adaptation dynamics), an aspect we have not studied in this paper.

Data and codes are available upon request.

## ACKNOWLEDGMENTS

We thank Jie Lin, Michael Manhart, Tzachi Pilpel and Yoav Ram for useful discussions and feedback. This research was supported by the National Science Foundation through the NSF CAREER 1752024 and the Harvard QBio fellowship.

## AUTHOR CONTRIBUTIONS

Y.G., A.A. designed research, performed research, and wrote the paper.

## Supplementary Information

### I. DETERMINISTIC GROWTH OF MUTANT FRACTION

Let *m* be the number of mutants with selection coefficient *s* in the population. If the mutant has established (which occurs with probability ≈ *s* for the Moran process), the dynamics of *m* can be approximated (in the large *N* limit) by the deterministic logistic equation:

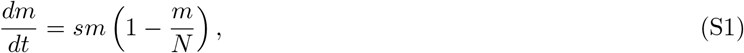

and its solution is given by:

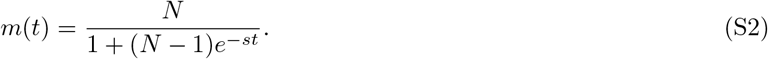

Defining the time *T* of the growth period as the time (in terms of the number of generations) to reach *N* − 1 mutants,

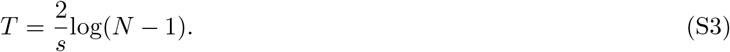

### II. FIXATION PROBABILITY OF A NEW MUTANT IN A POPULATION WITH TWO SUBGROUPS

Here we consider the scenario where there are two genotypes 1 and 2 with growth rates *g*_1_ and *g*_2_ = *g*_1_(1 + *s*) > *g*_1_ in a population of size *N* when a new mutant with growth rate *g_m_ = g*_1_(1 + *s_I_*) > *g*_1_ emerges.

Let *n* be the fraction of type 1 cells just before the mutant emerges. Assuming the dynamics follow a Moran process, and that no further mutations occur, the probability *p_ext_*[*n*] that the mutant goes extinct is the probability that this lineage goes extinct regardless of which event happens next:

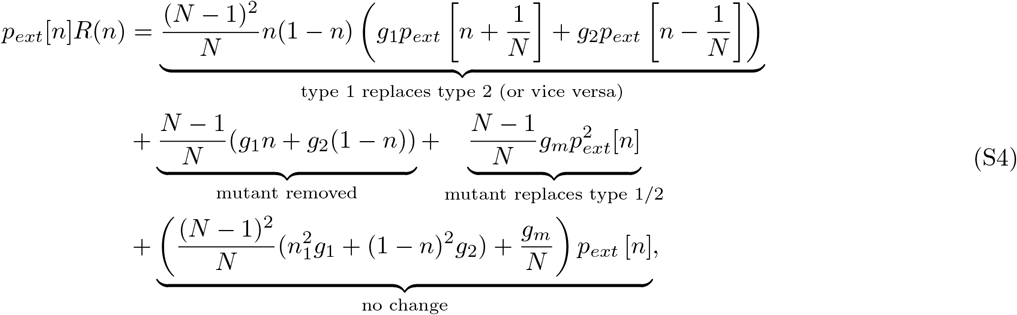

where the first term on the right takes into account the probability that the next event involves the division of a type 1 cell and the removal of a type 2 and vice versa, the second term involves the removal of the mutant, the third term is associated with the division of the mutant and removal of one of the other cell types, and the last term is for cases where there is no change in the composition of the population. *R*(*n*) is the total rate of all events and is the sum of the coefficients of all terms on the right of the equation:

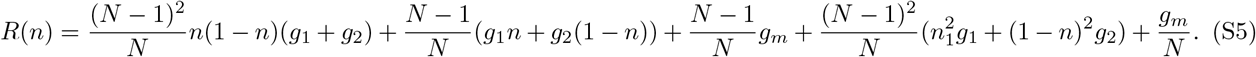

In Eqn. S4, we have also implicitly assumed that the population is large enough such that when there are two mutants, their fates are independent of one another.

Taking the large *N* limit, Eqn. S4 can be simplified to give:

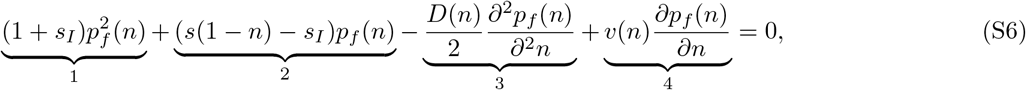

where 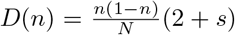, *v*(*n*) = *n*(1 − *n*)*s*, and the boundary conditions are given by *p_f_* (0) = (*s_I_ − s*)*/*(1 + *s_I_*) and *p_f_* (1) = *s_I_/*(1 + *s_I_*). In the large *N* limit, the third term will be much smaller than the other terms.

If the first two terms in Eqn. S6 dominate (either when *n* is close to 0 or 1, or when *s_I_* ≫ *s*),

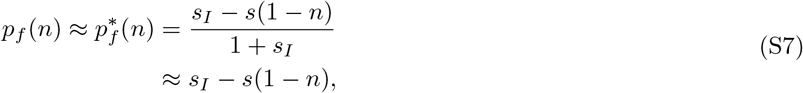

where in the second line we have taken the limit *s_I_* ≪ 1. For 0 < *n* < 1 and *s* > 0, the fourth term in Eqn. S6 is positive and hence this expression is an overestimate of the true fixation probability.

Denoting the actual fixation probability as

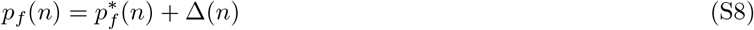

and substituting this back into Eqn. S6, we find that to first order in Δ and in the limit *s_I_* ≪ 1, Δ(*n*) satisfies the following equation:

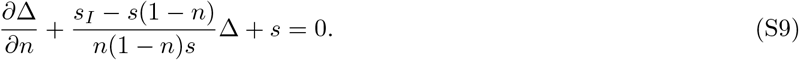

Defining the integrating factor:

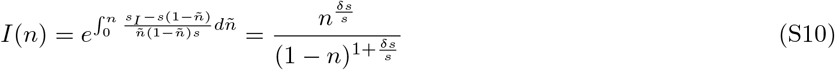

with *δs* = *s*^I^ − *s*, the solution to Eqn. S9 is given by:

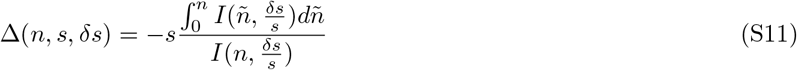

which satisfies the boundary conditions Δ(*n* = 0) = Δ(*n* = 1) = 0.

#### A. Comparison with exact values for the fixation probabilities

We find that these expressions for the fixation probability provide a good estimate of the exact values for a finite (but large) population (Fig. S1), which can be obtained as follows: Let 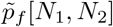 be the fixation probability of the mutant lineage when there are *N*_1_ type 1 cells and *N*_2_ type 2 cells in the population of size *N*, such that 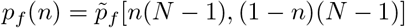. These fixation probabilities satisfy the following set of equations:

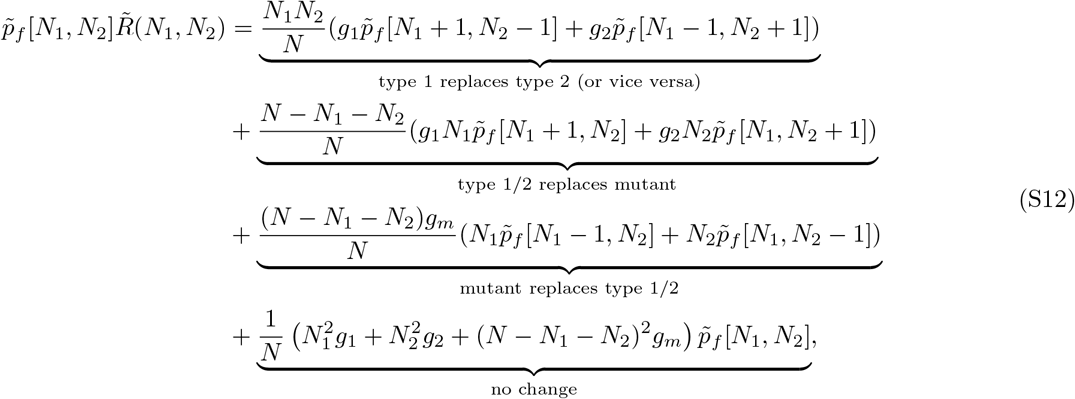

with boundary conditions 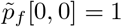 and 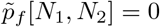 when *N*_1_ + *N*_2_ = *N*, and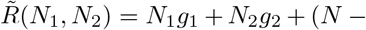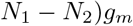 is the total growth rate of all cells.

This constitutes a large system of 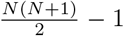 linear equations (Eqn. S12), that we solve by constructing a large matrix equation of the form 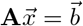 with 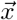 being the vector of fixation probabilites that we solve for.

**FIG. S1.**
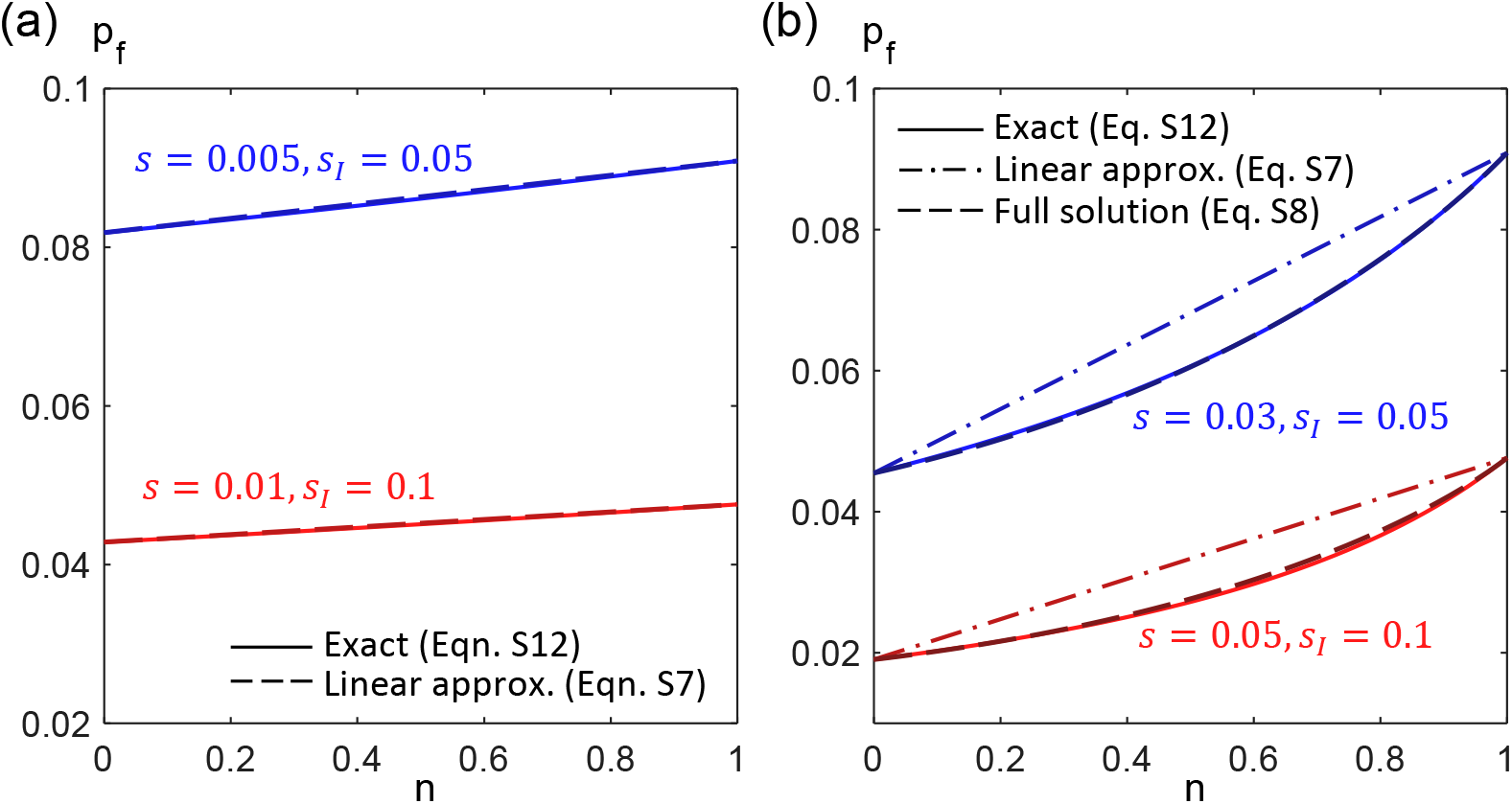
Fixation probability *p_f_* of a mutant (with growth rate *g_m_* = 1 + *s_I_*) as a function of *n*, the fraction of type 1 cells in a population of size *N* = 1000. Type 1 cells have growth rate *g*_1_ = 1 and type 2 cells have growth rate *g*_2_ = 1 + *s*. (a) When *s ≪ s_I_* (blue curves: *s* = 0.005, *s_I_* = 0.05, red curves: *s* = 0.01, *s_I_* = 0.1), *p_f_* (*n*) is approximately linear in *n* and is given by the selective advantage of the mutant over that of the population average. Solid lines are exact solutions obtained from Eqn. S12 while dashed lines plot the expression in Eqn. S7. (b) In general, *p_f_* (*n*) is not a linear function of *n* (dash-dotted lines, Eqn S7), but is well approximated by Eqns. S8, S11 (dashed lines vs. solid lines). [Blue curves: *s* = 0.03, *s_I_* = 0.05, red curves: *s* = 0.05, *s_I_* = 0.1.]

### III. EFFECT OF CLONAL INTERFERENCE ON FITNESS TRAJECTORIES

The probability that a successful interfering mutation emerges at time *t* after a mutation of fitness effect *s* has escaped loss via genetic drift and is still on its way to taking over the population is given by:

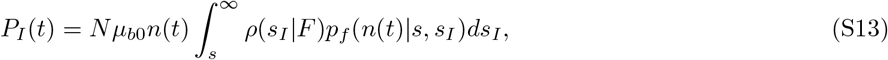

where 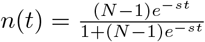 is the fraction of resident cells at time *t* (SI section I), *p_f_* (*n*(*t*)|*s, s_I_*) is the fixation probability of the interfering mutant with fitness effect *s_I_* (SI section II), and *ρ*(*s_I_ F*) is the distribution of fitness effects of beneficial mutations that can arise in a cell with fitness *F*.

Here, we consider a general class of exponential-like fitness effect distributions that has been adopted in previous theoretical studies [32, 33, 35]:

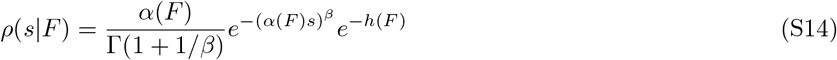

where Г is the Gamma function, *e*^−*h*^ is the fraction of mutations that are beneficial, and *α* and *β* characterize the shape of the distribution. In particular, if *β* = 1, we recover the exponential distribution used in the main text (Eqn. 2). When *β* > 1, *ρ*(*s*) falls faster than exponentially; when *β* < 1, *ρ*(*s*) falls more slowly than exponentially.

Substituting this distribution into Eqn. S13 gives:

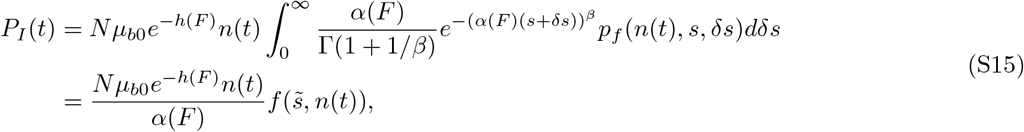

where 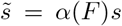, and *δs = s_I_* − *s*. For *β* = 1, if we assume that *p_f_* (*n*(*t*)*, s, δs*) = *δs + sn*(*t*) (Eqn. S7), then 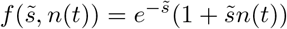.

Integrating this over all possible times the interfering mutant can emerge gives:

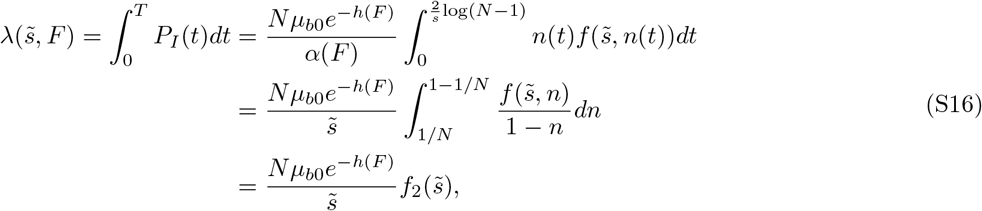

where 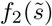 is a function of 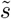. In the main text, we assumed the simple form of *p_f_* (*n*(*t*)*, s, δs*) = *δs + sn*(*t*) and *β* = 1, in which case 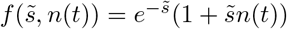 and hence

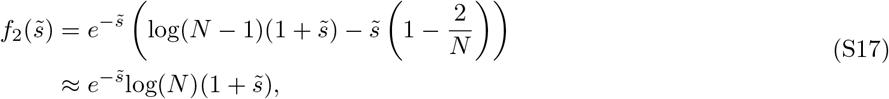

where we have taken the large *N* limit (log(*N*) ≫ 1)in the approximation. Substituting this into Eqn. S16, we recover Eqn. 10 in the main text.

Since the fixation probability *P_f_* is reduced by a factor of 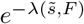 in the presence of clonal interference (Eqn. 5), the dynamics of average fitness is given by:

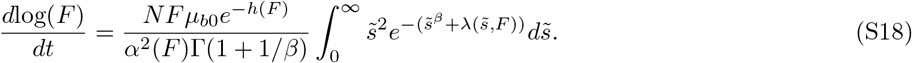

The fact that 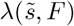 explicitly depends on *h*(*F*) but not *α*(*F*) implies that only *h*(*F*) affects the functional form of the fitness trajectory.

### IV. NUMERICAL RESULTS WITH OTHER DISTRIBUTIONS OF FITNESS EFFECTS

We find that our expression for the dynamics of fitness in the clonal interference regime (Eqn. S18) agrees well with simulation data for other distribution of fitness effects specified through different values of *β* (Eqn. S14), and that the conclusion that depletion in beneficial mutations speeds up the functional form of the trajectory holds for these different distributions (Fig. S2).

### V. SIMULATION DETAILS

We carry out full simulations of the evolutionary process within the Moran model using the Gillespie algorithm. For each simulation, the population is initialized with *N* cells, each with initial fitness (i.e. division rate) of 1. Throughout the simulation, we keep track of the size *N_i_* (i.e., number of cells) and fitness *F_i_* of cells in each subgroup *i* = 1, 2*, …, n_s_*, where *n_s_* is the number of currently existing subgroups that is present in the population. For example, the initial population consists only of *n_s_* = 1 subgroup of size *N*_1_ = *N* and fitness value *F*_1_ = 1. At each point during the simulation, we also calculate the corresponding beneficial mutation rates (i.e., probability of gaining a beneficial mutation per division) 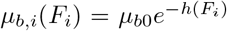 (Eqn. 2, Eqn. S14) of each subgroup. The possible events that change the composition of the population are: (a) a cell from a subgroup *i* divides (without mutation) and replaces a cell in another subgroup *j* /= *i*, which occurs with rate 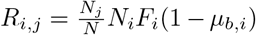, and (b) a cell from a subgroup *i* divides and mutates while a cell from subgroup *j* is removed which occurs with rate 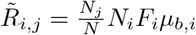 The total rate of an event occurring is then given by 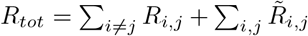 and we draw the time to the next event from an exponential distribution with mean given by 1*/R_tot_*. The next event is drawn with probability proportional to the rate at which the event occurs, and the composition of the population is updated accordingly. If a beneficial mutation occurs while a cell in subgroup *i* divides (i.e., event type b), we draw its selection coefficient from the specified distribution with mean given by 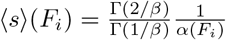 (Eqn. S14, or Eqn. 2 if *β* = 1), and this new mutant is stored as a new subgroup.

**FIG. S2.**
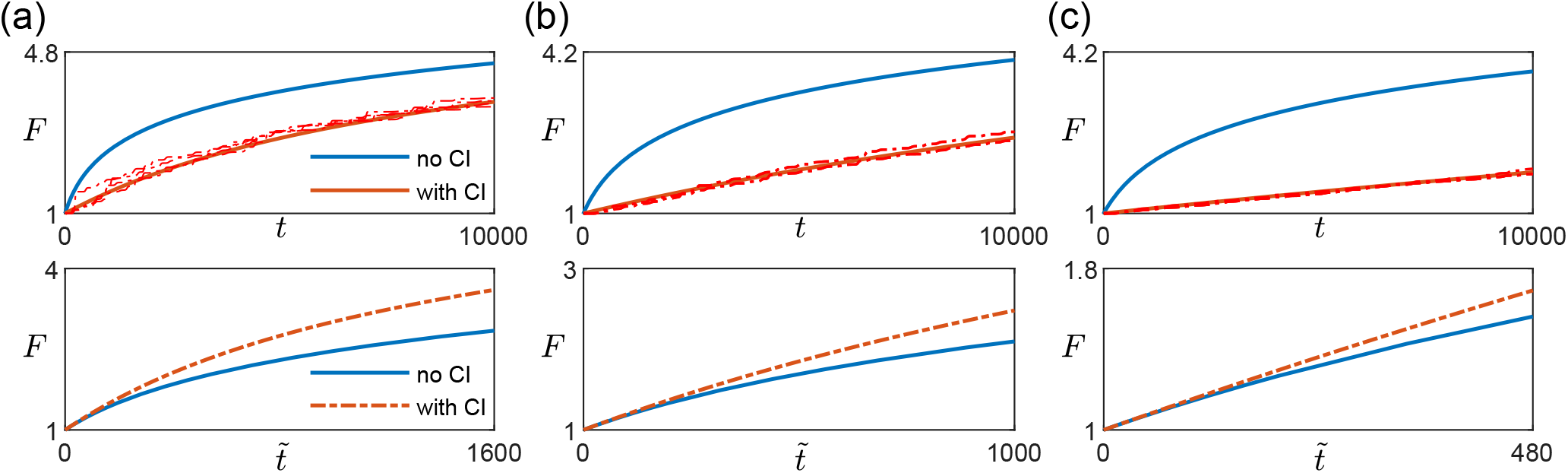
Fitness trajectories on the HoC/uncorrelated landscape with fitness effect distribution *ρ*(*s*) specified by (a) *β* = 0.5, (b) *β* = 1, and (c) *β* = 2. The parameter *β* characterizes how fast *ρ*(*s*) decays with *s* (Eqn. S14). In each sub-figure, the upper panel shows the fitness trajectories as a function of actual time with (red) and without (blue) clonal interference (CI). The solid lines are are obtained from solving Eqn. S18 (with *λ* given by Eqn. S16 in the CI regime and *λ* = 0 in the SSWM regime), while the dashed lines are individual replicates from simulations. The lower panel compares trajectories (blue and red solid lines in the upper panel) after rescaling time 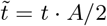 for the case with clonal interference such that both trajectories have the same derivative at *t* = 0. [Landscape parameters: 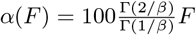, *μ*_*b*_ = *μ*_*b*0_*e*^−(*F* −1)^, other parameters: *N* = 10^6^, *μ*_*b*0_ = 1 *·* 10^−5^.]

### VI. EFFECT OF CLONAL INTERFERENCE ON SUBSTITUTION TRAJECTORIES

Analogous to the fitness trajectory, the average substitution trajectory *S*(*t*) (i.e. the number of fixed mutations as a function of time) is given by:

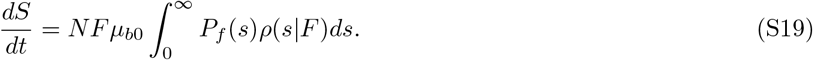

Assuming an exponential distribution of fitness effects (Eqn. 2), in the SSWM regime, the average substitution trajectory is therefore given by:

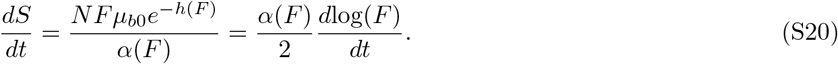

In the presence of weak clonal interference, assuming the simplified linear expression for the fixation probability (Eqn. S7) gives:

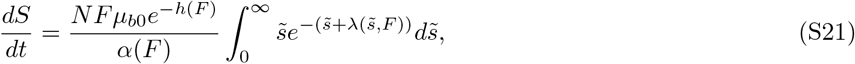

where 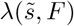 is the same as in the expression for fitness dynamics (Eqn. 10).

When comparing the numerical solutions of Eqns. S20 and S21, we scale time and *S* in the case with clonal interference such that 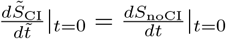

As for the fitness trajectory, clonal interference changes the form of the substitution trajectory only through *h*(*F*) (see Fig. S3 for an example). If the beneficial mutation rate *μ_b_* stays the same, the form of the substitution trajectory stays the same regardless of whether there is clonal interference. If *μ_b_* decreases with fitness, the form of the substitution trajectory speeds up; if instead *μ_b_* increases, the trajectory slows down.

**FIG. S3.**
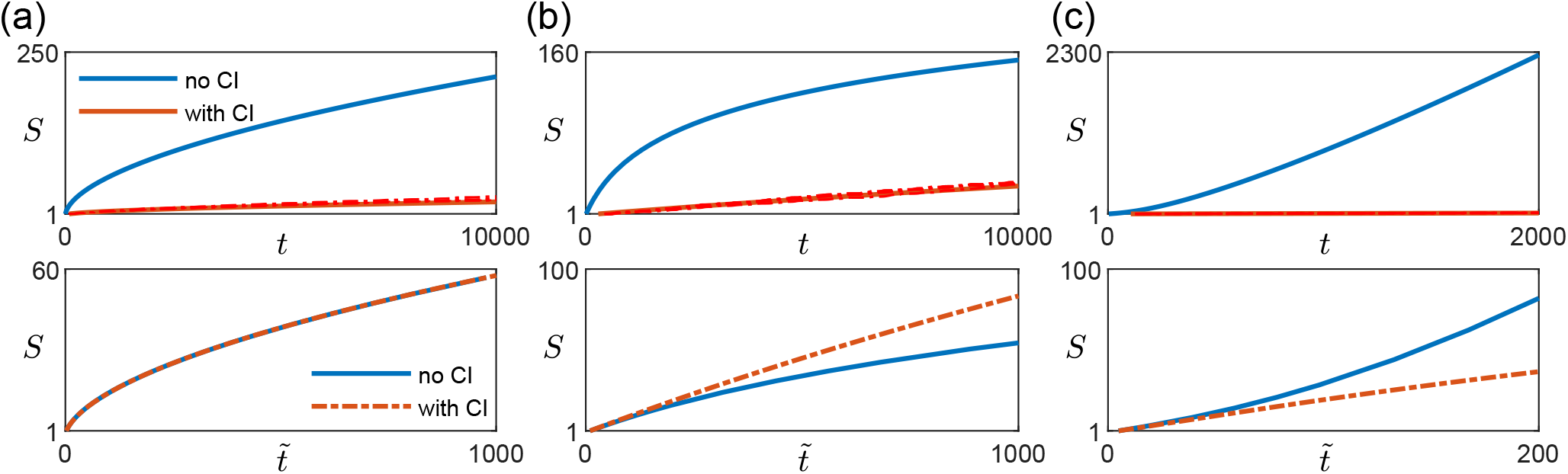
Effect of clonal interference on substitution trajectory for different types of macroscopic epistasis. (a) With the diminishing returns model where the mean fitness effect of beneficial mutations decreases with fitness but beneficial mutation rate stays the same, the functional form of the substitution trajectory stays the same. The upper panels show the substitution trajectories as a function of actual time with (red) and without (blue) clonal interference. The solid lines are are obtained from solving Eqns. S20 and S21, while the dashed lines are individual replicates from simulations. The lower panels compares trajectories (blue and red solid lines in the upper panel) after rescaling time 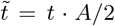 and 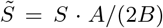 for the case with clonal interference such that both trajectories have the same derivative at *t* = 0. [Landscape parameters: *α*(*F*) = 20*F* ^5^, *μ_b_* = μ_*b*0_] (b) With the House-of-Cards/uncorrelated landscape where the mean fitness effects of beneficial mutations stays the same but the beneficial mutation rate decreases with fitness, the presence of clonal interference speeds up the functional form of the substitution trajectory. [Landscape parameters: *α*(*F*) = 100*F*, *μ*_*b*_ = *μ*_*b*0_*e*^−(*F* − 1)^] (c) With a mutation-releasing landscape where beneficial mutation rate increases with fitness, the presence of clonal interference slows down the functional form of the substitution trajectory. [Landscape parameters: *α*(*F*) = 50*e*^*F*−1^,*μ*_*b*_ = *μ*_*b*0_*e*^*F* − 1^; Other parameters: *N* = 10^6^, *μ*_*b*0_ = 1 *·* 10^−5^.]

